# Beyond the Hype: The Complexity of Automated Cell Type Annotations with GPT-4

**DOI:** 10.1101/2025.02.11.637659

**Authors:** Arman Kazmi, Deepshikha Singh, Shashank Jatav, Soumya Luthra

**Affiliations:** SoulBio, New Delhi

## Abstract

Recent research has shown the impressive capability of large language models like GPT-4 in various downstream tasks in single-cell data analysis. Among these tasks, cell type annotation remains particularly challenging, with researchers exploring various methods to improve accuracy and efficiency. While recent studies on GPT-like models have demonstrated annotation performance comparable to manual annotations, a significant gap remains in understanding their limitations and generalizability. In this work, we compare and evaluate the annotation performance of the GPT-4 model against traditional methods on nine randomly selected public single-cell RNA seq datasets from cellxgene, covering diverse tissue types. Our evaluation highlights the complexity of annotating cell types in single-cell data, revealing key differences between automated and manual approaches. We found specific cases where GPT-4 underperforms, demonstrating its limitations in certain contexts. We further introduce an automated approach to incorporate literature search using a RAG approach which enhances and outperforms GPT-4 cell type annotation when compared to traditional methods. We also introduce metrics based on taxonomic distance in the ontology tree to evaluate the granularity of the cell type annotations. To support future research, we also release an open-source Python package ^1^ that enables fully automated cell-type annotation of single-cell data using GPT-4 alongside other methods. The pipeline can take paper as an input and do cell type annotations on its own.

## 1 Introduction

The recent research in large language models (LLMs) like GPT-4 has demonstrated impressive potential in biomedical research. Their ability to process complex and high dimensional data makes them particularly suitable for applications like Single-Cell RNA Sequencing (scRNA-seq) analysis. This technique has been used to study cellular heterogeneity by profiling hundreds and thousands of individual cells. It also helps in identifying novel cell types and states, uncovering cellular interactions, and characterizing the microenvironment (Quan et al., 2023).

One of the important steps in scRNA-seq data analysis is cell type annotation, which helps in understanding the specific functions of different cell types within tissues. The accurate annotation of cell types is very important for understanding various disease mechanisms, particularly in conditions like cancer and autoimmune disorders where cellular composition and interactions play a critical role. Traditionally, cell-type annotation has been a manual process that is both labor-intensive and technically demanding. It requires biologists to meticulously compare the highly expressed genes within each cell cluster against established canonical marker genes to accurately identify cell types. To address these challenges, various automated cell type annotation approaches have been developed. In general, automated approaches can be categorized into three types: (1) marker gene-based annotations, (2) correlation-based annotations, and (3) supervised classification-based annotations (Pasquini et al., 2021).

Marker gene-based annotations is one of the widely used approaches where the typical “cluster then annotate” paradigm is followed (Cao et al., 2019). The clusters are derived from unsupervised learning and then annotated based on the matching of cluster-level gene expression to predefined marker gene lists (Pasquini et al., 2021). CellMarker (Zhang et al., 2019b) and PanglaoDB (Franzén et al., 2019) databases act as central repositories, which systematically index thousands of marker genes across human and mouse cell types from diverse studies. Tools like SCType (Ianevski et al., 2022) use these databases to compare gene expression profiles with known cell type markers. They generate similarity scores to enable accurate and reliable cell type annotations. Although marker-based annotations are quite popular they have certain limitations as well such as the selection of marker genes is heavily dependent on prior knowledge, which can introduce bias and errors (Pasquini et al., 2021). Also, not all cell types have well-defined marker genes particularly novel or rare cell types which often lack established markers. This is mostly true in the case of complex datasets, where cell types are often characterized by multiple genes rather than single markers.

Correlation-based methods compare gene expression profiles of clusters to established reference datasets to determine the most closely matching cell types (Pasquini et al., 2021). Tools like SingleR (Aran et al., 2019) and scmap (Kiselev et al., 2018) use reference data like the Human Primary Cell Atlas (Mabbott et al., 2013) to annotate individual cells without predefined clusters which enhances flexibility for heterogeneous datasets. Although effective for datasets with well-characterized reference data, these methods are constrained by the quality, completeness, and potential biases of the reference data, often resulting in suboptimal performance for rare or novel cell populations.

Supervised classification-based methods are based on the classical machine learning approach where patterns from gene expression profiles are learned from labeled data (training set) and used to predict cell types in unlabeled data (test set) (Pasquini et al., 2021). Tools like CellAssign (Zhang et al., 2019a) incorporate prior knowledge of marker genes into probabilistic models to assign cell types, while scGPT (Cui et al., 2024) uses transformer-based architectures pre-trained on large annotated datasets to classify cells. These methods enhance accuracy and scalability but their limitations lie in their reliance on the quality and representativeness of labeled data, which can affect accuracy and generalizability, especially in cases of cellular heterogeneity from mixed or overlapping cell types.

Despite these advancements, limitations in traditional tools highlight the need for more adaptable and accurate approaches. There has been a recent surge in deep learning-based methods for scRNA-seq analyses claiming superior performance by addressing the limitations inherent in traditional supervised based-methods (Wang et al., 2019; Menden et al., 2020; Wang et al., 2021, 2022; Dong et al., 2024). Deep learning models, including LLMs like GPT-4, offer a novel approach by integrating extensive biological knowledge with computational adaptability. The introduction of transformer-based models particularly the BERT (Devlin et al., 2019) architecture has revolutionized the field of Natural Language Processing (NLP) with remarkable progress due to the powerful self-attention mechanism and its ability to process and integrate long-range information. Following the BERT architecture, the scBERT (Wang et al., 2022) model, represents a breakthrough in single-cell RNA-seq analyses. Trained on large-scale unlabeled scRNA-seq data, scBERT effectively addresses batch effects and enhances generalizability in cell type annotations.

More recently, advancements in high-performance computing have enabled the development of Large Language Models (LLMs) like GPT-4 (Achiam et al., 2023) and LLaMA (Touvron et al., 2023). These LLMs have a unique ability for transfer learning and can leverage knowledge learned from the general domain to develop expertise in specific areas, often performing better than models trained only on domain-specific data. Recent studies (Hou and Ji, 2024) have shown that GPT-4 achieves remarkable accuracy in cell-type annotation, with over 75% concordance with manual annotations across diverse datasets. Tools such as GPTCelltype, designed to interface with GPT-4, streamline its integration into workflows like Seurat, avoiding the need for additional pipelines or extensive reference datasets. The authors used marker genes for each cluster and experimented with different prompt strategies to improve cell type annotations. For evaluations, the authors used a scoring system assigning values of 1, 0.5, or 0 to “fully match,” “partial match,” and “mismatch” outcomes, respectively, comparing GPT-4’s annotations to manual annotations. Average scores across cell types and tissues were calculated to determine accuracy. However, partial match scoring does not inform about the distance between GPT-4’s prediction and manual annotation. This lack of information limits our ability to assess the granularity or specificity of cell type annotation.

In this work, we build on the work of Hou and Ji, expanding the evaluation of GPT-4’s annotation capabilities using marker genes across 9 datasets randomly selected from CELLxGENE (Abdulla et al., 2023) platform. We first obtain the top differential genes using SCANPY (Wolf et al., 2018) and then use a basic prompt with GPT-4 to annotate the cell types. We then normalized the obtained annotations in the cell ontology using SapBERT (Liu et al., 2021) to evaluate the automated annotations. For example, if GPT-4 predicted T cell type, we map it to its standard term in cell ontology. We further compare the GPT-4’s annotation performance against traditional approaches, including SingleR (Aran et al., 2019), CellMarker2.0 (Hu et al., 2022), ScTypeDB, and ScType with Cell MarkerDB (Ianevski et al., 2022), using the numeric scoring method outlined in Hou and Ji. We further expand the evaluation criteria by introducing a taxonomic distance-based scoring metrics which are based on the distance between the automated annotations and the manual annotations in the ontology tree to evaluate the granularity or specificity of the automated annotations.

In our analysis, we found that while the GPT-4 annotations align with the manual annotations in most of the cases, our findings highlight specific cases where it underperforms. To address this, we employ a chain-of-thought prompting strategy enriched with context retrieved from the published literature of the datasets using a retrieval agent. This approach significantly improves the annotations by leveraging contextual information from the published literature, enabling annotations that align more closely with manual annotations and improving the reproducibility of author-defined cell types. Further, to support future research on cell type annotation we release a Python package for GPT-4 cell type annotations along with other traditional methods.

## 2 Data

In this study, we selected nine diverse single cell RNA sequencing datasets (scRNA-seq) from the CELLxGENE platform (Abdulla et al., 2023) to evaluate the annotation capabilities of GPT-4. CELLxGENE, an open source tool developed by the Chan Zuckerberg Initiative, provides a repository of curated single cell datasets across various tissues and species, making it a valuable resource for large-scale single-cell analysis. This platform enables researchers to access a wide array of datasets with standardized metadata, interactive visualization tools, and expression profiles at the single-cell level.

We chose data sets representing a range of tissue types: prostate, blood, retina, fallopian tube, lung, heart, colon, gastric, esophagus, and brain. These datasets encompass both well-characterized and complex tissues, providing a balanced representation of cell type diversity and heterogeneity. Each dataset was retrieved using a unique dataset identifier from CELLxGENE, ensuring consistent and reproducible data collection. Table 1 details all the datasets used in this study for evaluating the annotation capabilities of GPT-4. The datasets include cell counts ranging from approximately 17,000 to 100,000, capturing a wide range of cell type abundance and variability. This heterogeneity, combined with the inclusion of both normal and disease states such as benign prostatic hyperplasia, HIV infectious disease, Barrett esophagus, and opiate dependence, ensures a comprehensive evaluation across diverse biological contexts. The selection of the datasets allows for a robust and comprehensive assessment of GPT-4’s annotation capability and testing its generalizability across various tissue types and disease states. Further, the heterogeneity of the cells in the datasets represents real-world scRNA-seq data analysis scenarios that require annotation methods to perform well on both abundant and rare cell populations. This diversity ensures that the evaluation captures the complexity of biological systems, including subtle differences in cell states and identities.

**Table 1:**
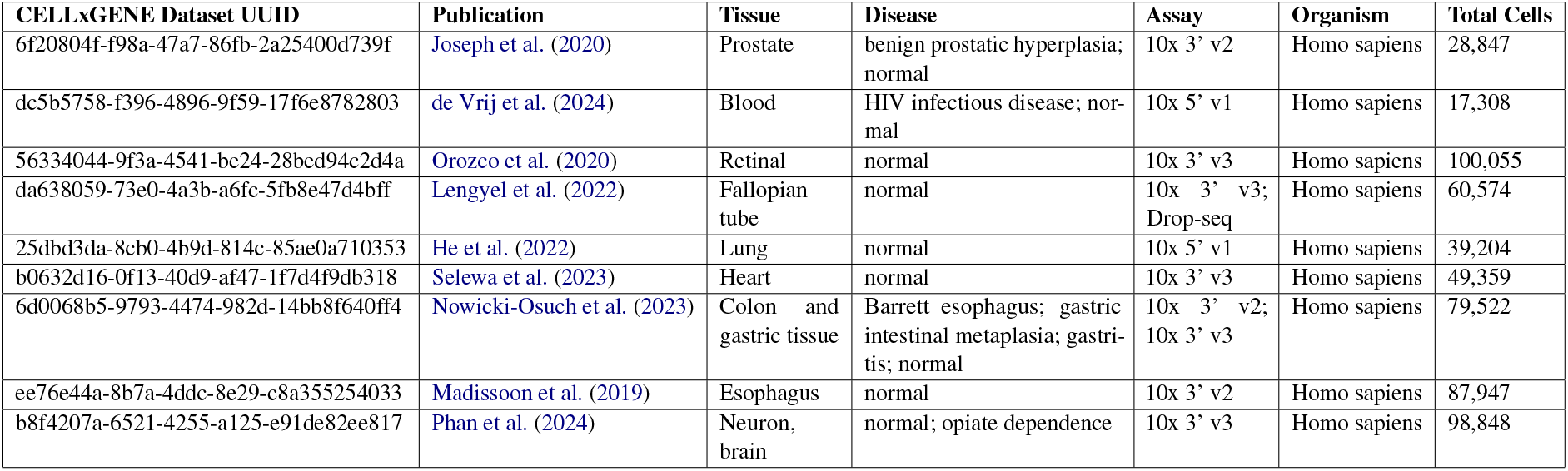
CELLxGENE dataset details.

## 3 Methods

In this section, we describe in detail the method and experiments for annotating the cell types.

### 3.1 Cell type annotation methods

Since the goal of this paper is to evaluate GPT-4 capabilities of cell-type annotations as claimed in Hou and Ji (2023), we explain in detail the cell type annotation method using GPT-4 alongside the other popular traditional approaches followed in this work. All the implementation were done in Python 3.11.

#### 3.1.1 GPT-4

We first ran SCANPY (Wolf et al., 2018), a Python tool for scRNA-seq analysis, on each dataset to find the top differentially expressed genes for each cluster. It performs a t-test on the gene expression data, grouping cells based on their Leiden cluster assignments. For each cluster, the top 10 and top 20 differentially expressed genes were extracted based on their ranking.

For annotations, we used the GPT-4 - 13 June 2023 version, the same as the version used in the (Hou and Ji, 2023) to make the comparison fair. After getting the top differentially expressed we use the basic prompt template which takes the top marker genes for annotations. The basic prompt template used was same as that of Hou and Ji:

*Identify cell types of TissueName cells using the following markers separately for each row. Only provide the cell type name. Do not show numbers before the name. Some can be a mixture of multiple cell types. \n Marker Gene List*.

#### 3.1.2 SingleR

SingleR Aran et al. (2019) employs a reference-based approach for annotation by comparing the gene expression profiles of individual cells to those of a reference dataset. This method generates a score for each potential cell type to assign the most appropriate annotation based on the highest scores. In this work, We used the Python implementation of SingleR to get the “best” cell type for each cluster. For the reference dataset, we used the Human Primary Cell Atlas (HPCA)^2^ and then calculated the dominant cell type of each cluster. To identify dominant cell types for each cluster, we calculated the frequency of cell types within each cluster based on “best” cell types. For clusters with three or more cell types, we identified cell types comprising at least 50% of the cluster as dominant. The three most frequent cell types were selected if no single cell type met this threshold. For clusters with fewer than three cell types, all were included.

#### 3.1.3 CellMarker2.0

We used the CellMarker 2.0 (Hu et al., 2022) web tool-CellMarker annotation platform for cell type annotation. The top differentially expressed genes from each cluster, identified through our single-cell RNA-seq analysis, were provides as input to the tool. CellMarker 2.0 matched these genes to its curated tissue and disease-specific marker genes database for accurate annotations of the cell types. The results were used to assess the accuracy of CellMarker 2.0 annotations by comparing them with the original cell types.

#### 3.1.4 ScTypeDB

ScType (Ianevski et al., 2022) is a popular machine-learning-based classification algorithm that uses both supervised and unsupervised learning techniques to identify and classify cell types. This method generates a score based on the similarity of gene expression profiles to known cell types, enabling accurate annotations. We used the Python implementations of SCTypeDB. The gene expression data was processed by converting gene names to uppercase and selecting genes present in predefined gene sets (gs and gs2) from ScTypeDB. To convert cell-level scores to cluster-level scores, we aggregated the scores of individual cells within each cluster, summing the scores for marker genes and sorting them to identify the top-ranking cell types. We took the top-scoring cell type as the annotation for that cluster.

#### 3.1.5 ScType with Cell MarkerDB

For an alternative approach, cell type annotation was performed using CellMarkerDB ^3^-a comprehensive database of marker genes for various human cell types.

### 3.2 Cell type evaluation

To compare the annotations generated by GPT-4 and other methods with manual annotations, we first normalized the outputs from each method. For normalization, we utilized SapBERT (Liu et al., 2021) with the Cell Ontology^4^. SapBERT requires a dictionary containing the names and synonyms of the ontology’s terms. It uses a self-alignment mechanism to generate dense embeddings for biomedical terms, enabling accurate mapping of annotations to their corresponding concepts in the preferred ontology.

We first follow the matching criteria mentioned in (Hou and Ji, 2023) to evaluate the normalized outputs against the manual annotations. In particular, we classify the normalized annotation as ‘fully match’ if normalized and manual annotations are the same, and ‘partial match’ if the normalized annotation is either the parent or child of the manual annotation in the cell ontology tree. Finally ‘mismatch’ if the normalized annotation differs from the manual annotation and is not in the same lineage in the cell ontology. For numeric comparison, the score assigned is 1, 0.5, and 0 for fully, partial, and mismatch respectively, and similar to the paper we calculate the average score within each dataset across cell types. To extend the evaluation framework, we incorporate additional metrics that assess the granularity and depth of partial matches, as these metrics provide critical insights into the granularity or specificity of the annotations in the ontology tree. The following metrics are defined:

- **Average Score**: This metric represents the mean score across all clusters in a dataset and is defined as:

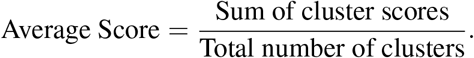 It serves as an overall indicator of annotation performance.
- **Average Taxonomic Distance**: For partial matches, the taxonomic distance represents the distance in the ontology tree between the normalized and manual annotations. The average taxonomic distance is computed as:

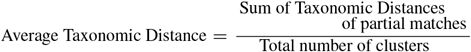 For cases where there are no partial matches the average taxonomic distance would be 0. A lower average taxonomic distance means the automated annotations are more closer to the manual annotations in the cell ontology tree.
- **Granularity Index**: This metric combines the average score and the Taxonomic Distances to assess both the accuracy and proximity of annotations, defined as:

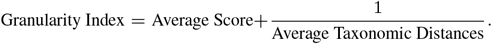 A higher granularity index indicates closer and more accurate matches, whereas a lower index reflects distant or mismatched annotations. For cases when there is no partial matches the granularity index is not defined.

### 3.3 Improving annotations using Retrieval

To improve the annotation quality of the GPT-4 we employ a chain-of-thought prompt strategy with adding additional contexts from the published literature of the datasets. For every dataset, we refer to their publication and retrieve context to use with the chain-of-thought prompting using Langchain ^5^ and GPT-4. We use the URL of the published literature of the dataset and query the GPT-4 to extract the marker gene information for the particular tissue of the dataset and then use that information in the chain-of-thought prompting to annotate the clusters.

The query template which was used to create the retrieval agent was:

*Extract all the mentioned cell types and marker genes (if any) from the paper for TissueName*.

The chain-of-thought prompt template that was then used with retrieval was:

*You are an expert in cell type annotation for TissueName. Your task is to identify cell types based on the provided marker genes for each cluster. Please follow these guidelines: \n 1. Analyze each cluster separately and provide a specific cell type name for each. \n 2. Some clusters may represent a mixture of multiple cell types or unknown cell types. Be precise in your annotations. \n 3. Use the given marker genes for each cluster to determine the cell type. \n 4. Additional context from the research paper is also provided below. Use this information to refine your annotations, but do not limit yourself to only these cell types. \n 5. Integrate the provided context with your broader knowledge of cell types and markers to make the most accurate and precise annotations possible. \n 6. If a cluster’s identity is ambiguous or doesn’t match known cell types, provide your best estimate or label it as ‘Unknown’. \n MarkerGene List \n Additional Context retrieved from the paper*.

## 4 Results

Our analysis evaluated the performance of a large language model (LLM)-based cell type annotation applied to single-cell RNA-sequencing (scRNA-seq) datasets from various tissues. We assessed the GPT-4 model’s predictions both with and without a retrieval-based grounding step, comparing them against established manual annotations and known cell type ontologies. Across multiple datasets, including prostate, human blood, fallopian tube, heart, colon, gastric, esophagus, neuron/brain, and human retinal tissue, the model’s retrieval-enabled predictions demonstrated improved concordance with expert-assigned cell types and established ontologies.

### 4.1 Cell Type Annotations without Retrieval

Here we report the annotations result when GPT-4 was queried only using basic prompting and with the top 10 and 20 marker genes. Table 2 shows the average scores of the GPT-4 annotations with basic prompting compared to the other traditional methods. The highest score is highlighted in each row. There is a slight difference in the performance of GPT-4 when the number of marker genes is increased from the top 10 to the top 20 as shown in Fig 3. The average scores remain nearly equal or show a slight improvement with the top 20 marker genes, except in the case of the Nowicki-Osuch et al. (2023) dataset, where GPT-4 did not perform as expected, with an average score of only 0.08, possibly due to overfitting to less relevant genes and also the dataset contains relatively rarer cell types. Among traditional methods, CellMarker 2.0 performs the worst, with an average score of 0 in most cases, possibly due to limited marker database and lack of context specific understanding while SingleR performed comparatively better, possibly due to larger marker database. In three cases, SingleR matches GPT-4’s performance in one case and outperforms it in the other two cases.

**Table 2:**
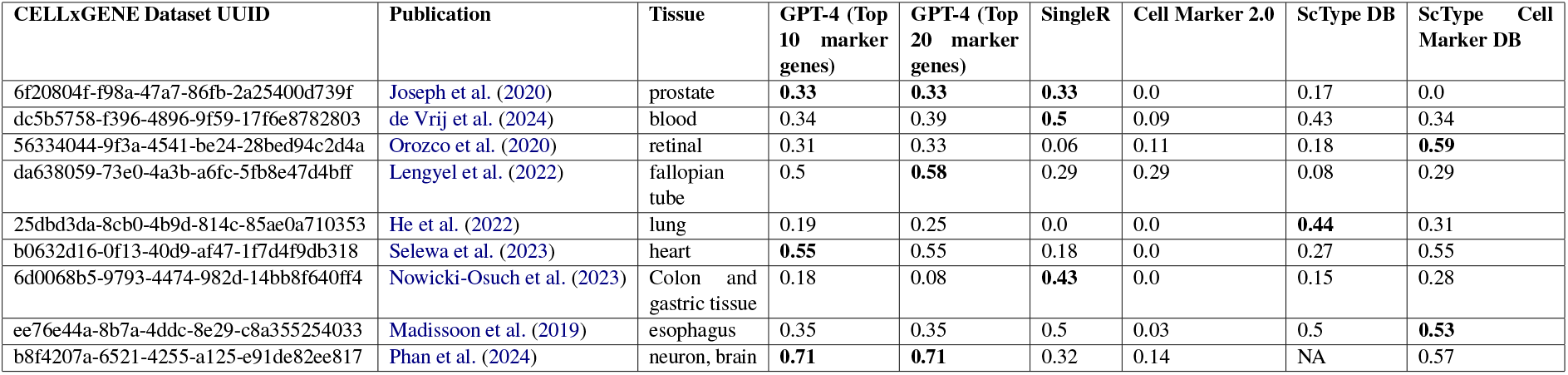
Average scores of annotation methods across different datasets and tissues without retrieval.

It is interesting to note that methods other than GPT-4 do not give consistent performance in all the datasets. For example, SingleR does not perform great in Heart, Retinal, Lung and Neuron tissues while SCtypeDB does not perform great in Prostate, Fallopian tube, Retinal, Colon and Gastric and Neuron tissues. SCtype CellMarkerDB does not perform good in Fallopian tube, Prostate and Neuron tissues, while CellMarker 2.0 does not perform well in any of the tissues. This performance drop is most likely due to the limitations of datasets these methods are trained on. For example, we used “HPCA” datasets for SingleR which does not have a lot of Fibroblast cells in the training data while it has abundant cells from Blood tissue. This explains SingleR’s good performance in Blood and poor performance in Lung.

### 4.2 Cell Type Annotations with Retrieval

When additional context was given to GPT-4 with a chain-of-thought prompting strategy, GPT -4 annotation improved performance. Fig 3 shows the average scores of GPT-4 annotations with and without retrieval on CELLxGENE datasetes when queried using the top 10 and the top 20 genes. The performance increased in most of the cases when the top 10 marker genes was used as input. On dataset Nowicki-Osuch et al. (2023) where GPT-4 has an average score of 0.18 increased to 0.38 when retrieval was used with the top 10 marker genes. Fig 1 shows the partial matches of GPT-4 in case of retrieval and without retrieval with the top 10 and top 20 marker genes. The dashed line on above of each bar category represents the total number of clusters in that particular dataset.

**Figure 1:**
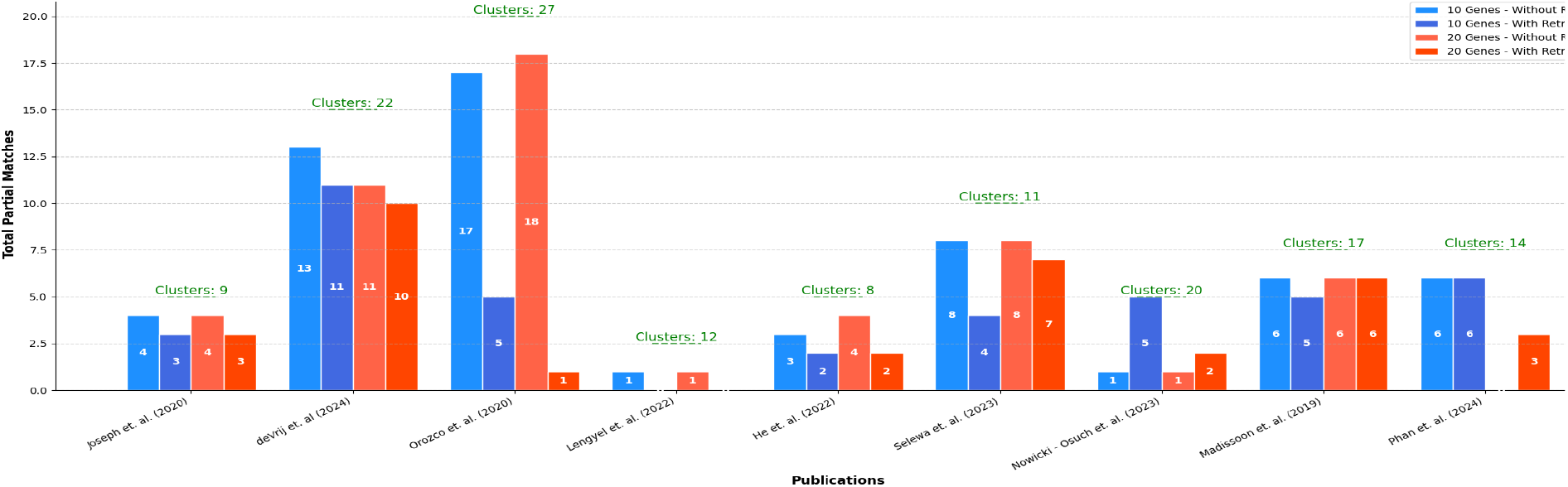
Comparison of Total Partial Matches With and Without Retrieval for Top 10 and 20 Marker Genes. Dashed lines represent the number of clusters for each dataset. Without retrieval the partial matches are greater than those with retrieval in most of the cases

**Figure 2:**
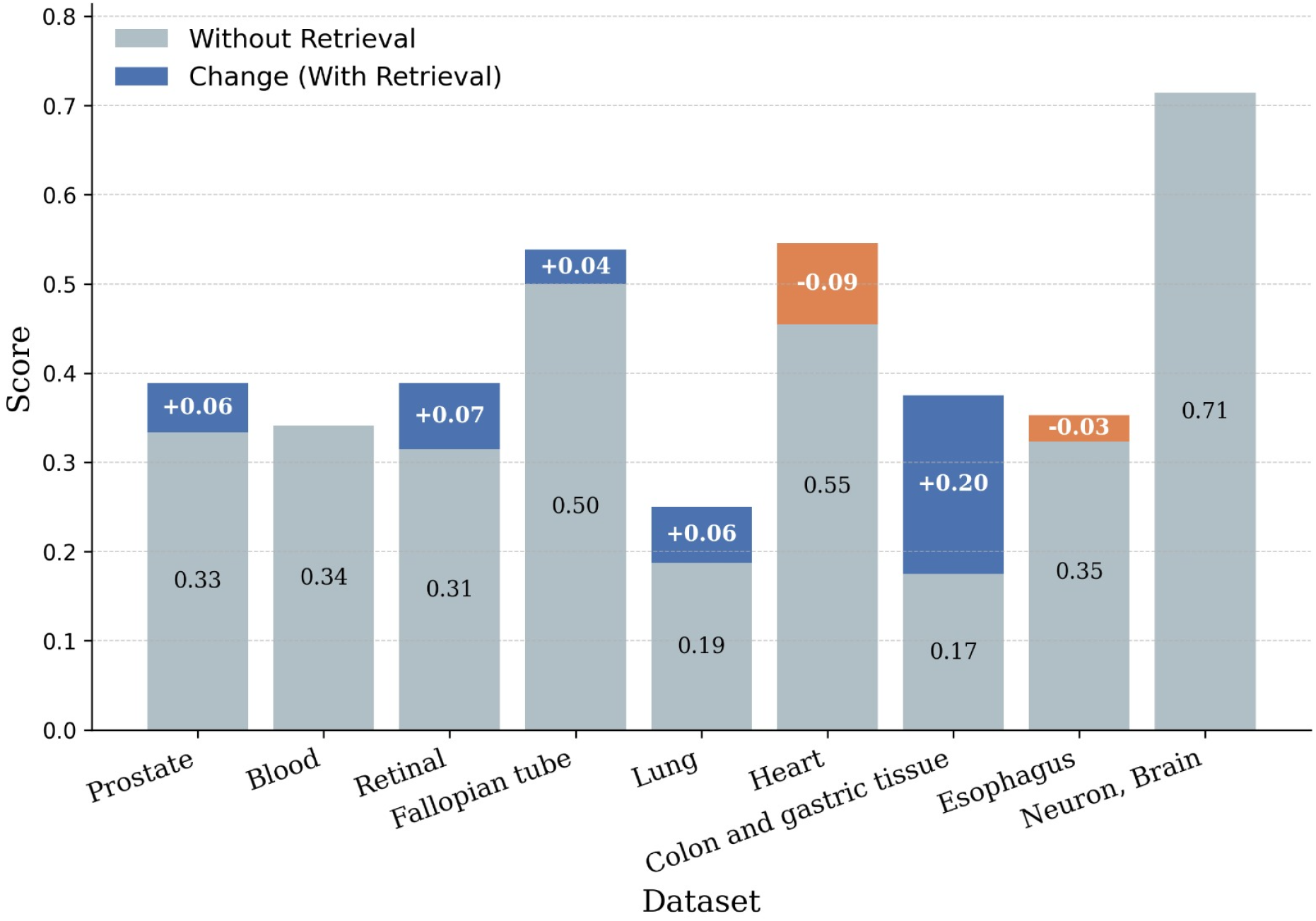
Comparison of Average Scores With and Without Retrieval for Top 10 Marker Genes.

**Figure 3:**
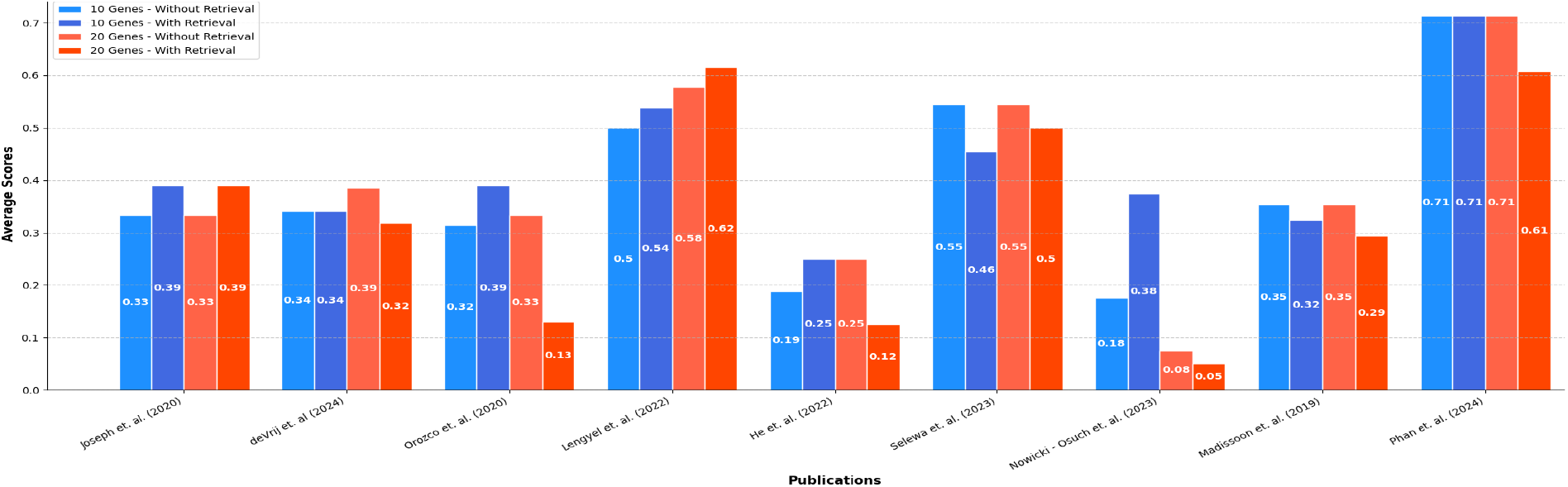
Comparison of Average Scores With and Without Retrieval for Top 10 and 20 Marker Genes.

Overall, the retrieval-augmented model exhibited a marked increase in accuracy for well-defined cell types that have distinct transcriptomic signatures. For example, canonical immune cell types such as B cells, T cells, macrophages, and monocytes were consistently identified correctly. In the human blood dataset (de Vrij et al., 2024), clusters characterized by genes like TYROBP, LYZ, and FCN1 were correctly identified as monocytes by the retrieval approach, aligning with the manual annotation of “CD14-positive monocyte.” Similarly, endothelial cells and smooth muscle cells in tissues like the fallopian tube (Lengyel et al., 2022) and the heart (Selewa et al., 2023) were recognized with high accuracy, with retrieval frequently achieving a full or partial match with manually assigned identities. Notably, the retrieval step helped the model navigate complexities in epithelial subpopulations. In gastrointestinal tissues, such as those from the colon and stomach (Nowicki-Osuch et al., 2023), retrieval-assisted predictions identified specialized epithelial subsets (e.g., enterocytes, mucous neck cells, and foveolar cells) with greater precision. While both retrieval and non-retrieval predictions often captured the broad cell class (e.g., “Epithelial Cells”), the retrieval-based method was more likely to refine this categorization to the appropriate cell subtype. For instance, the cluster characterized by CDH17, PRSS3, and GPA33 was predicted as “Enterocytes” or “Epithelial Cells” in the retrieval mode, showing closer alignment with known intestinal absorption cell identities compared to the generic epithelial predictions made without retrieval. In neural tissues (Phan et al., 2024), retrieval allowed the model to differentiate more nuanced cell classes. Oligodendrocytes, astrocytes, oligodendrocyte precursor cells, and microglia were identified in a manner consistent with manual annotations. Retrieval-based predictions reduced confusion between closely related cell types, such as differentiating oligodendrocytes from oligodendrocyte precursor cells, and microglia from macrophage-like populations. Similarly, in the human retinal (Orozco et al., 2020), the retrieval mode generally supported more accurate identification of well-known retinal subtypes (e.g., amacrine cells and bipolar neurons), although some specialized subtypes (e.g., rod versus cone photoreceptors) remained challenging in certain clusters. Despite these improvements, some inconsistencies persisted. In complex cases, where gene expression signatures overlapped across cell types or represented poorly characterized subpopulations, the model-even with retrieval-defaulted to more generic cell type identities. For example, certain stromal or fibroblast-like cells in lung or prostate tissues were occasionally misclassified as other cell types, reflecting the inherent ambiguity in their transcriptomic profiles. Similarly, some specialized gastric cell types, such as parietal cells or enteric endocrine lineages, still required further refinement. While retrieval-based predictions were more accurate than those without retrieval, they occasionally mixed these subtypes or assigned overly broad identities.

It is interesting to note that that without retrieval the partial matches are greater than those with retrieval in most of the cases as in Fig 1. The retreival cases has lower number of partial matches and more exact matches which is why the average score are greater than without retrieval in most of the cases.

## 5 Discussion

Quantitatively, retrieval reduced Average Taxonomic Distance and increased the Granularity Index across most tissues Table 3, with the greatest improvements in Retinal (0.185 vs. 1.444 Avg. Tax. Dist., 5.794 vs. 1.007 GI), Lung, and Heart. Prostate and Blood showed moderate gains, while the Fallopian tube achieved a perfect match. Colon or Gastric Tissue and Esophagus had minor increases in taxonomic distance but maintained reasonable granularity. These results highlight retrieval as a powerful tool for refining annotation accuracy.

**Table 3:** Average taxonomic distance and granularity index for GPT-4 annotations (Top 10 genes) with and without retrieval. The best score for each datasets in each category are highlighted in bold.

| Dataset UUID | Publication | Tissue | Avg. Tax. Dist. |  | Granularity Index |  |
| --- | --- | --- | --- | --- | --- | --- |
|  |  |  | With Retrieval | Without Retrieval | With Retrieval | Without Retrieval |
| 6f20804f-f98a-47a7-86fb-2a25400d739f | Joseph et al. | Prostate | **1.111** | 1.444 | **1.29** | 1.026 |
| dc5b5758-f396-4896-9f59-17f6e8782803 | de Vrij et al. | Blood | **1.1818** | 1.4091 | **1.187** | 1.0505 |
| 56334044-9f3a-4541-be24-28bed94c2d4a | Orozco et al. | Retinal | **0.185** | 1.444 | **5.794** | 1.007 |
| da638059-73e0-4a3b-a6fc-5fb8e47d4bff | Lengyel et al. | Fallopian tube | 0 | 0.083 | NA | 12.54 |
| 25dbd3da-8cb0-4b9d-814c-85ae0a710353 | He et al. | Lung | **0.25** | 0.375 | **4.25** | 2.85 |
| b0632d16-0f13-40d9-af47-1f7d4f9db318 | Selewa et al. | Heart | **0.4545** | 0.81 | **2.654** | 1.779 |
| 6d0068b5-9793-4474-982d-14bb8f640ff4 | Nowicki-Osuch et al. | Colon and gastric tissue | 0.4 | **0.05** | 2.875 | **20.175** |
| ee76e44a-8b7a-4ddc-8e29-c8a355254033 | Madissoon et al. | Esophagus | 1.05 | **0.647** | 1.275 | **1.898** |
| b8f4207a-6521-4255-a125-e91de82ee817 | Phan et al. | Neuron, brain | **0.785** | **0.785** | **1.987** | **1.987** |

The incorporation of a retrieval-based grounding step significantly enhanced the LLM’s ability to accurately identify and classify cell types from scRNA-seq data. By utilizing curated ontologies and context from literature searches, the retrieval approach provided the model with reference points against which to anchor its predictions, leading to improved accuracy over the without-retrieval approach. This benefit was particularly pronounced in cell types with well-established marker sets and distinct functional roles such as monocytes, B cells, endothelial cells, and astrocytes where the retrieval guidance reinforced the identification of canonical gene signatures.

A key advantage of retrieval lies in its completely automated process and capacity to resolve ambiguities. Without retrieval, the LLM tended to default to generic categories or closely related cell classes, particularly in heterogeneous tissues where multiple cell types share overlapping gene expression patterns. Retrieval contextualized the GPT-4 reasoning, allowing it to differentiate between, for example, “endothelial cell” and “endothelial cell of lymphatic vessel,” or “smooth muscle cell” versus “fibroblast,” based on known and defined hierarchies present in established ontologies.

Nonetheless, the observed limitations indicate that improvements are still needed. The retrieval step, while improves with the help of additional context but still wasn’t able to achieve annotation performance at par with manual annotations. More complex or less canonical cell types with incomplete or evolving marker definitions remained challenging. Additionally, for rare or less-characterized subsets, retrieval might provide limited improvement if reliable references are scarce or ambiguous. From a broader perspective, the retrieval-enhanced approach underscores the importance of integrating structured biomedical ontologies and knowledge bases with LLM-based inference methods. As curated cellular identity ontologies (e.g., Cell Ontology) and comprehensive gene-marker databases expand, we can expect further refinements in cell annotation accuracy. Moreover, iterative improvements such as incorporating user feedback, adopting more specialized ontologies, and continuously updating the retrieval corpus could enable the model to navigate the subtle distinctions within complex cellular landscapes.

A major application of all these methods is to reproduce already published literature along with their raw data since most of the times authors fail to provide cell annotations for the datasets. So in these cases it’s known which cell types are being dealt with beforehand, which makes it possible for any researcher to make an informed method of choice for annotating cell types. The way to do that could be to first annotate broader cell types using GPT-4 along with retrieval and then use traditional methods like SCtype trained on tissue of interest since performance of traditional methods depends on datasets they are trained on.

## 6 Conclusion

In this study, we investigated the performance of GPT-4 in cell-type annotation using marker genes with different prompting strategies. We compared its results to traditional methods such as SingleR, Cell Marker 2.0, ScType DB, and ScType with Cell Marker DB. Our evaluation, which focused on annotation accuracy, average taxonomic distance and granularity concordance with manual annotations, revealed several key insights into the capabilities and limitations of GPT-4 and traditional methods. We further integrated contexts from literature search of the datasets to assess impact on annotation performance and found that GPT-4 performance does improve when it’s guided with relevant context. Our study shows that GPT-4 is not a one-size-fits-all-solution for cell type annotation but it is consistent and gives average performance in all contexts. Traditional methods specifically trained on specific tissues and cell-types will outperform GPT-4 but we can only know of target tissues while reproducing a paper, so GPT-4 might be a better choice when annotating cells for the first time.

In conclusion, the retrieval-based LLM predictions represent a substantial step toward more reliable and fine-grained automatic cell annotation. These findings highlight not only the feasibility of applying LLMs in a biomedical context but also emphasize the pivotal role of knowledge-grounding methodologies. With continued improvements in both computational models and reference databases, retrieval-augmented LLMs hold considerable choice for accelerating and improving the standardization of single-cell dataset annotations across diverse tissues and species.

While GPT-4 does perform better at cell type annotation in most of the case it is important to consider it’s limitations as mentioned in Hou and Ji. Some of the major concerns while using GPT-4 for annotation cell types is the corpus it has been trained on which is unknown. Another limitation is the potential for bias in its predictions, as GPT-4 relies on patterns in the data it has seen rather than direct experimental validation. Additionally such large model may struggle with rare or novel cell types that are underrepresented in its training corpus and may hallucinate. Therefore, while GPT-4 can be a valuable tool for assisting in cell type annotation, its predictions should be interpreted with caution and validated by a domain expert.

Since the release of GPT-4, the version used in this work, there has been significant development in GPT-4-like models. These models have been shown to possess reasoning capabilities such as o1 ^6^, which may improve the performance of cell type annotations, as they can reason before providing a final annotation. Another potential improvement could be fine-tuning GPT-4 or any foundation model on a high-quality reference dataset.

## Footnotes

1 https://github.com/soulbio/cell_type_annotation

2 https://www.humancellatlas.org

3 http://117.50.127.228/CellMarker/CellMarker_download.html

4 https://www.ebi.ac.uk/ols4/ontologies/cl

5 https://python.langchain.com/v0.1/docs/

6 https://openai.com/o1/

